# Extensive Expansion of the *Speedy* gene Family in Homininae and Functional Differentiation in Humans

**DOI:** 10.1101/354886

**Authors:** Liang Wang, Hui Wang, Hongmei Wang, Yuhui Zhao, Xiaojun Liu, Gary Wong, Qinong Ye, Xiaoqin Xia, George F. Gao, Shan Gao

## Abstract

**Background:** The cell cycle plays important roles in physiology and disease. The *Speedy*/RINGO family of atypical cyclins regulates the cell cycle. However, the origin, evolution and function of the *Speedy* family are not completely understood. Understanding the origins and evolution of *Speedy* family would shed lights on the evolution of complexity of cell cycles in eukaryotes.

**Results:** Here, we performed a comprehensive identification of *Speedy* genes in 258 eukaryotic species and found that the *Speedy* subfamily E was extensively expanded in Homininae, characterized by emergence of a low-Spy1-identify domain. Furthermore, the *Speedy* gene family show functional differentiation in humans and have a distinct expression pattern, different regulation network and co-expressed gene networks associated with cell cycle and various signaling pathways. Expression levels of the *Speedy* gene family are prognostic biomarkers among different cancer types.

**Conclusions:** Overall, we present a comprehensive view of the *Speedy* genes and highlight their potential function.

## Background

Cell cycle progression is tightly controlled via periodical activation of the cyclin-dependent kinases (CDKs)[1], which are modulated by cyclin binding and subsequent phosphorylation of an evolutionarily conserved threonine in the T-loop of the CDKs[2]. *Speedy/RINGO* (herein referred to as *Speedy*) is a novel CDK regulator, which was first identified as able to induce rapid maturation of oocytes in *Xenopus laevis* by mediating the G2/M transition[3]. Although it has no sequence homology to any known cyclins, *Speedy* is able to fully activate CDKs by binding to their *Spyl* domains[4, 5]. This process is also independent of CDK threonine phosphorylation[6, 7]. Numerous *Speedy* genes have been identified in mammals with different expression profiles and CDK binding preferences.

*Speedy* A, C and E1 can bind to and activate Cdk1, Cdk2, or Cdk5 with different preferences to regulate cell cycle progression, and they have different expression profile[8, 9]. Under physiological conditions, *Speedy* A is also a requisite for Cdk2 targeting to telomeres in mice and can induce G2/M transition [10], but *Speedy* E1 impairs this progression in *Xenopus*[11]. It has been shown that deregulation of *Speedy* genes is associated with tumor. For instance, high *Speedy* A expression levels mediate the initiation and progression of tumorigenesis in breast cancer[12]. *Speedy* A impedes the functional differentiation of growable neural stem/progenitor cells in clonally derived neurospheres in glioma, and simultaneously increase their number and longevity[13]. Further, its expression level also correlates with tumor grade and poor survival in gliomas. Taken together, *Speedy* genes play an important role in both normal tissues and tumors.

Thus far, the evolution of *Speedy* genes has been studied in 12 species of vertebrates, and seven *Speedy* genes have been identified in the human genome[14]. However, the *Speedy* family is not completely understood from the context of evolution, and the *Speedy* family nomenclature is inconsistent among different studies. With advances in new sequencing techniques and the important function of the *Speedy* family in cell cycle and cancer, it is important to comprehensively identify *Speedy* members in all eukaryotic species and study their potential roles in human tissues and tumors. Such information will shed light on the evolution of complexity of cell cycles in eukaryotes.

In this study, we performed a systematic identification and phylogenetic analysis of *Speedy* family genes. Like cyclins, the *Speedy* family also tends to expand in higher animals. Particularly *Speedy* subfamily E extensively expands in Homininae, accompanied by the emergence of a novel domain architecture, and they are neither conserved within Homininae nor fixed in the human population. Further functional studies also indicate they have distinct CDK preferences and effects on the cell cycle. Additionally, their expression profiles imply their potential function in both human brain development and tumors. Finally, we propose that genes generated by duplications (paralogs) might form a competitive endogenous RNA (ceRNA) network with its parental and cognate genes.

## Methods

### Sequence Retrieval

We used 2 protein sequences (NP_877433.2 and XP_004342966.1 from *Homo sapiens* and *Capsaspora owczarzaki* ATCC 30864, respectively) as seeds to search against NCBI databases, including nonredundant protein database, EST database, genome database and transcriptome database, by BLAST. As known, Speedy proteins so far only contained the *Spy1* domain, so we used this criteria to identify other potential *Speedy* family members in this study. Sequences which contained only *spy1* domain, screening by interproscan [15, 16], were retained. To obtain more complete datasets, we used the *spy1* domain HMM model from Pfam [17] to search against the proteome of genome-available species which have Speedy proteins, by using HMMER v3.1b2 [18]. Coding sequences were then extracted from their corresponding genomic or transcript sequences by using BLAT [19] and Genewise [20]. For genome-available species, only the longest transcript of each gene was retained for further analysis. After removing redundant sequences for each species, the retained coding sequences were then used for further analysis.

### Phylogenetic and Population Analysis

Sequences from 15 genome available species (*Danio rerio*, *Mus musculus*, *Homo sapiens*, *Bos mutus*, *Gallus gallus*, *Xenopus Silurana tropicalis*, *Ursus maritimus*, *Loxodonta africana*, *Oryzias latipes*, *Ailuropoda melanoleuca*, *Myotis lucifugus*, *Ciona intestinalis*, *Chelonia mydas*, *Vicugna pacos*, *Calypte anna*, using *Amoebidium parasiticum* JAP-7-2 as the outgroup) were selected to reconstruct the phylogeny of the *Speedy* family. Protein sequences were aligned by PROBCONS v1.12[21], with default parameter except for the option of iterative refinement, for which we used 1000 iterations. We then backed the alignment to its corresponding coding sequences. After getting conserved blocks from sequence alignment by Gblocks v0.91b [22, 23], we used jModelTest v2.1.6 [24, 25] to find the best substitution model according to Bayesian Information Criterion. Afterwards, PhyML 20141106[26] were used to reconstruct phylogenetic tree under the best substitution model 012212+G+I, in which gamma was 1.32 and proportion invariable sites was 0.13, with bootstrapping of 1000 replicates. WebLogo was used to visualize the sequence alignment[27].

Sequences of the *Speedy* family from species belonging to Catarrhini were extracted. We found that the length of them varied wildly, which was not suitable for further analysis. Sequences meeting the following criteria were removed from further analysis: a *spy1* domain length shorter than 100aa; if the start site of *spy1* domain was at the beginning of sequence, which is shorter than 180aa. We then used the same protocol described above to reconstruct the phylogenetic tree by using cds sequences. The best model for phylogenetic tree reconstruction was 012010+G (with gamma = 3.59). Type ◻ and ◻ diversity of the *Speedy* family were detected by DIVERGE 3.0 Beta 1[28, 29]. PAML 4.8 [30, 31] was used to detect positive selection for *Speedy* E family in Catarrhini with branch-site model and calculate the dN/dS ratio for low-Spy1-identity domain sequences. Sequence alignment were visualized by WebLogo[27]. Coding potential score for human and chimpanzee *Speedy* members were obtained from the Coding Potential Calculator[32].

SNP data in cds and cdna region of genes and pseudogenes of *Speedy* family were extracted from the 1000 Genomes Project Phase 3 data [33]. For protein-coding genes of the human *Speedy* family, SNPs which could be a result of the stop codon were extracted, and if simultaneous SNPs around adjacent sites led to a stop codon, individual data were then used to determine whether there were simultaneous mutations within these sites in each individual. As for pseudogenes, we focused on SNPs which could change a stop codon in the middle of coding region to a code for an amino acid. We also extract CNV data in genes and pseudogenes of the *Speedy* family to study whether copy number variation existed in these regions. Fst was calculated by VCFtools v0.1.14-30[34]. Data process was performed with scripts, written in Java.

### Gene expression analysis, survival analysis and co-expression network construction

The RNASeq data of 32 human normal tissues were downloaded from Human Protein Atlas Project[35]. Transcriptomic datasets of human prefrontal cortex were from [36]. Only one sample for each tissue was randomly selected to further analysis. Bam files of 734 cancer cell lines from 15 cancer types were downloaded from the CCLE project[37]. For normal tissue data, all reads were mapped to the human genome (GRCh37 in Gencode v25) by HISAT2 [38]. FPKM was calculated by Stringtie [39] for both human normal tissues and cancer cell lines data. We then calculated the tissue specificity of *Speedy* members using the method described in [40]. We also calculated the correlation of expression profile for each pair between *Speedy* members (expressed in more than one sample). We used a Pearson correlation coefficient >=0.8 and adjust *p* value <0.05 (Bonferroni correction) as criteria to determine whether a pair of *Speedy* members had similar expression profiles. Days-to-last-followup and days-to-death representing the number of days from initial pathological diagnosis to the last time the patient was known to be alive or dead, respectively, were extracted from TCGA clinical information and used to construct Kaplan-Meier survival curves by the R survMisc package[41]. To find a combination of gene signatures to predict survival with the expression of Speedy members, we first used the univariate Cox proportional-hazards model to eliminate genes, in which the absolute z score was less than 2. According to this criteria, all 7 members were retained. The multivariate Cox proportional-hazards regression model was used to select suitable combination of genes with backwards elimination algorithm by MASS package in R[42]. In the final model, we calculated risk score for each patient by sum up the product of log transformed FPKM and its corresponding coefficient from multivariate Cox proportional-hazards regression model. The formula was as follows: risk score = 0.553\**SPDYA*+0.195\**SPDYE4*+0.314\**SPDYE6*-0.516\**SPDYE7P*+0.364\**SPDYE8P*. WGCNA[43] was used to construct weighted gene co-expression networks for normal tissues and cancer cell lines. We first manually removed those low expressed genes (FPKM<0.00001 in more than 90% samples). FPKM was then log-transformed by using log2(x+1). After sample clustering, outlier samples, (if they exist), were also removed from datasets. Retained genes and samples were used to calculate soft power (the smallest threshold making scale free topology with R >=0.9, which could result in a good balance between scale-free fitness, mean of connectivity and modularity). This threshold was then used to calculate the topological overlap matrix (known as TOM). The soft power for normal tissues was 3 and datasets of each kind of cancer type were listed in Table S6.

ClueGO 2.3.2[44] and CluePedia 1.3.2[45] were used to functional enrichment analysis and visualization of genes which co-expressed with *Speedy* members. As co-expressed genes varied wildly among different *Speedy* members, we used different criteria to deal with this problem. When compared the functional enrichment of co-expressed genes in normal tissues, the parameter “min genes” were 7, 5 and 3 for SPDYE4, SPDYE1 and SPDYA, respectively. Parameter “%genes” were 8%, 6% and 4% for SPDYE4, SPDYE1 and SPDYA, respectively. When compared to the functional enrichment of co-expressed genes of SPDYE4 between normal tissues and HNSC, the same parameters were used. We used GO term fusion parameter to reduce the redundancy. Only Biological Process were used in our functional enrichment analysis.

### MiRNA target prediction and ceRNA networks construction

We used 3 different microRNA target prediction tools to predict microRNA targets for 3’UTR of protein-coding genes and cdna of pseudogenes in *Speedy* family using human microRNA in miRBase release 21 as references. TargetScan[46], PITA[47] and miRanda[48] (with -strict parameter) were used in this study. Targets shared by all 3 methods were then considered to be the most credible. Gene pairs with correlation coefficient of expression profiles >=0.8 with adjust pvalue < 0.05 (Bonferroni correction) and shared same microRNAs were considered to be function as ceRNA to each other. ceRNAs networks were visualization by Cytoscape 3.3[49].

## Results

### The distribution and evolution of *Speedy* family genes

To comprehensively study the evolution of the *Speedy* family, we first identified all *Speedy* family protein-coding genes. According to our search strategy described in the methods, we found 544 *Speedy* genes in 258 eukaryotic species (Fig. 1a and Table S1). These species belong to 11 phyla, 30 classes, 112 orders, 195 families, and 237 genera, respectively. All of the species belonged to Metazoa except for four species, which belonged to Ichthyosporea (*Amoebidium parasiticum* JAP-7-2 and *Capsaspora owczarzaki* ATCC 30864) and Amoebozoa (*Dictyostelium fasciculatum* and *Dictyostelium purpureum*), respectively. In Metazoa, the Mammalia, Aves, Clupeocephala, and Protostomia have more *Speedy* genes than other classes. Interestingly, zebrafish contain *Speedy* genes, whereas carp fish do not. A *Speedy* gene is also found in the jewel wasp genome, but is absent in the bee and fruit fly genomes. Overall, these data indicate that *Speedy* genes minimally originated before the common ancestor of Metazoa, and they are unevenly distributed across species.

**Figure 1.**
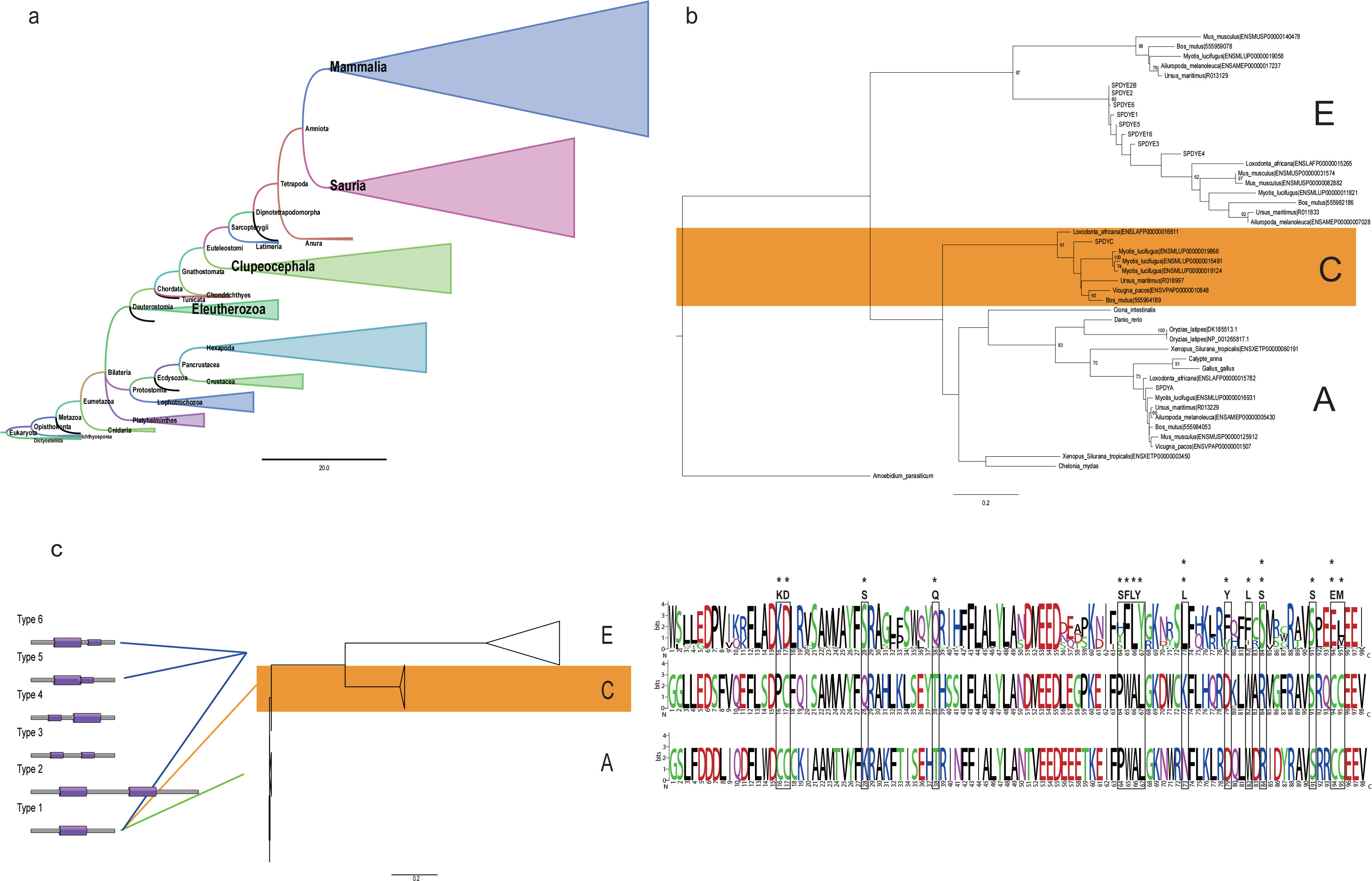
A comprehensive landscape of the *Speedy* family. (a) Distribution of *Speedy* genes among species. The size of the triangles is proportional to the number of species within each clade. (b) Phylogenetic tree of the *Speedy* family. The *Speedy* family can be split into three subfamilies named A, C, and E. (c) Schematic domain architecture of *Speedy* genes (left panel) and sites under positive selection for subfamily E (right panel). The conserved residues of *Spy1* domains are displayed in sequence logos among the Catarrhini. The relative size of the amino acid letters in the logos represents the raw frequency for the alignment in each subfamily. The positive sites are indicated by asterisks using Bayes Empirical Bayes (* > 0.95; ** > 0.99).

We also found that the number of *Speedy* genes varied among species (Table S1). More than half of the investigated species (148 of 258) contained only one *Speedy* gene. These species were mainly non-mammals (144 out of 148). In mammals, most species (76 out of 80) contained more than one *Speedy* gene. For instance, Catarrhini species contained at least four *Speedy* genes, and the human genome contains 10 *Speedy* protein-coding genes. Taken together, these data demonstrate that higher primate species possess more *Speedy* genes.

To explore the evolutionary relationships among *Speedy* genes, we constructed a phylogenetic tree with the conserved region of the *spy1* domain of 15 representative species, using the maximum-likelihood method and *A. parasiticum* JAP-7-2 being as the outgroup. According to the phylogenetic tree, *Speedy* genes were divided into three subfamilies, designated as subfamilies A, C, and E following the name of the human *Speedy* gene in each clade (Fig. 1b). Subfamily A exists in all selected species, suggesting that this subfamily originated first, while subfamilies C and E originated after A. Apparently, subfamily C was lost in mice, further supporting the uneven distribution of the *Speedy* family across species. Extensive expansion of *Speedy* subfamily E with a novel domain architecture was found in Homininae

In addition to the uneven distribution of *Speedy* genes across species, we also found that two distinct domains exist among Speedy proteins using InterProScan[15, 16]. The major domain is a full-length *Spy1* domain (e value ≤ 1e^−24^), but we also found that there is a relatively low-Spy1-identity domain in some proteins (e value ~ 1e^−15^) after screening. We further focused on the domain architecture of Speedy proteins and their distribution across species. Out of 544 proteins, 508 contained a full *Spy1* domain, while others contained two *spy1* domains and could be further divided into five additional categories based on the identity and location of the *Spy1* domains (Fig. 1c). *Balaenoptera acutorostrata scammoni* and *Papio anubis* contain a protein with two full *Spy1* domains, respectively. *Ornithorhynchus anatinus* and *X. laevis* contain a protein with two low-identity *Spy1* domains. The remaining organisms contain both a full-Spy1 and a low-identity *Spy1* domains. Furthermore, 30 of 32 were Catarrhini species, and all of their low-identity domains are C-terminal, (named LSI-C). The others belong to *Myotis davidii* and *Tupaia chinensis* and are N-terminal. We identified one, seven, and 13 LSI-Cs genes in the gorilla, human, and chimpanzee genomes, respectively, and also found that most syntenic regions of human LSI-C genes were gaps in gorillas, indicating that the number of LSI-Cs in gorillas could be underestimated due to the quality of the gorilla genome (Table S2). Clearly, LSI-Cs are new genes that recently originated in the common ancestor of Catarrhini and then extensively expanded in Homininae. According to the phylogenetic analysis, all LPI-Cs from Catarrhini were assigned to subfamily E (Fig. 1c).

We further examined whether the low-Spy1-identity domain was functional or only displayed sequence similarity. After calculating the rate of nonsynonymous to synonymous nucleotide mutations (dN/dS) among low-Spy1-identity domain sequences across the Catarrhini, we found that > 90% of the dN/dS ratios were < 1 (Fig. S1a), suggesting that they are subject to negative selection. This indicates that the low-*Spy1*-identity domain is functional.

We also investigated if the full-length *Spy1* domains in subfamily E are undergoing positive selection. We used PAML and found that 15 amino acids (aa) in the *Spy1* domain of subfamily E display positive selection. Among them, 12aa have posterior probabilities ≥0.95, and the others were ≥0.99 in the branch-site model test[30, 31] (Fig. 1c).

Furthermore, we also wondered whether the copies in subfamily E have divergent functions compared to those in subfamilies A and C. The coefficient of functional divergence (θ) was used to determine the type ◻ divergence (functional changes) between them by DIVERGE 3.0[28, 29]. The θ for C to E and A to E were 0.662391 and 0.561062, respectively, which are both significantly >0 (Z-score test, *p* <0.01), indicating that subfamily E has functionally differentiated. Together, these results suggest that the genes in subfamily E have functionally differentiated from those in subfamilies A and C.

### *Speedy* subfamily E in the human genome

Next, we focused on the *Speedy* family in humans. In addition to 10 protein coding genes, we also identified 14 *Speedy* pseudogenes in the human genome, all of which were paralogs of four LSI-Cs, and nine of them were paralogs of the *SPDYE3* gene (Fig. S1b)[50]. Taking the 10 protein-coding genes into account, there are 24 *Speedy* members in the human genome. When we examined the locations of these genes, we found that most of subfamily E members are located on chromosome 7 (Fig. 2a), except *SPDYE4* and *SPDYE22P*. We next sought to determine if human *Speedy* members exist in other Homininae. We found that 21 human *Speedy* members exist in the chimpanzee genome, but most of them (14 out of 24) were lost in gorillas (Fig. 2b). Then, we calculated the coding potential value (CPV) of the sequences in the Homininae, which is positively correlated with the protein coding capacity of a given sequence calculated by the Coding Potential Calculator[32] to examine whether they had protein-coding capacity. For subfamily C, the CPVs are nearly the same, but that of *SPDYA* in gorillas is much lower. When we compared the CPVs of the protein coding genes in subfamily E between the human and chimpanzee genomes, we found that counterparts in chimpanzees had significantly lower CPVs than in humans (paired sample sign test, *p* = 0.00531, Fig. S1c). Moreover, stop codons are sparse in the counterparts of human coding DNA sequences (CDSs) except for *SPDYE3*, *SPDYE4*, *SPDYE5*, and *SPDYE16* in chimpanzees and *SPDYE1*, *SPDYE4*, *SPDYE5*, and *SPDYE16* in gorillas, indicating that protein coding capacity was lost for some *Speedy* members in both chimpanzees and gorillas. The same trends were also found in pseudogenes (*p* = 0.005). Taken together, these data suggest that subfamily E members are not conserved in Homininae, except for *SPDYE4*.

**Figure 2.**
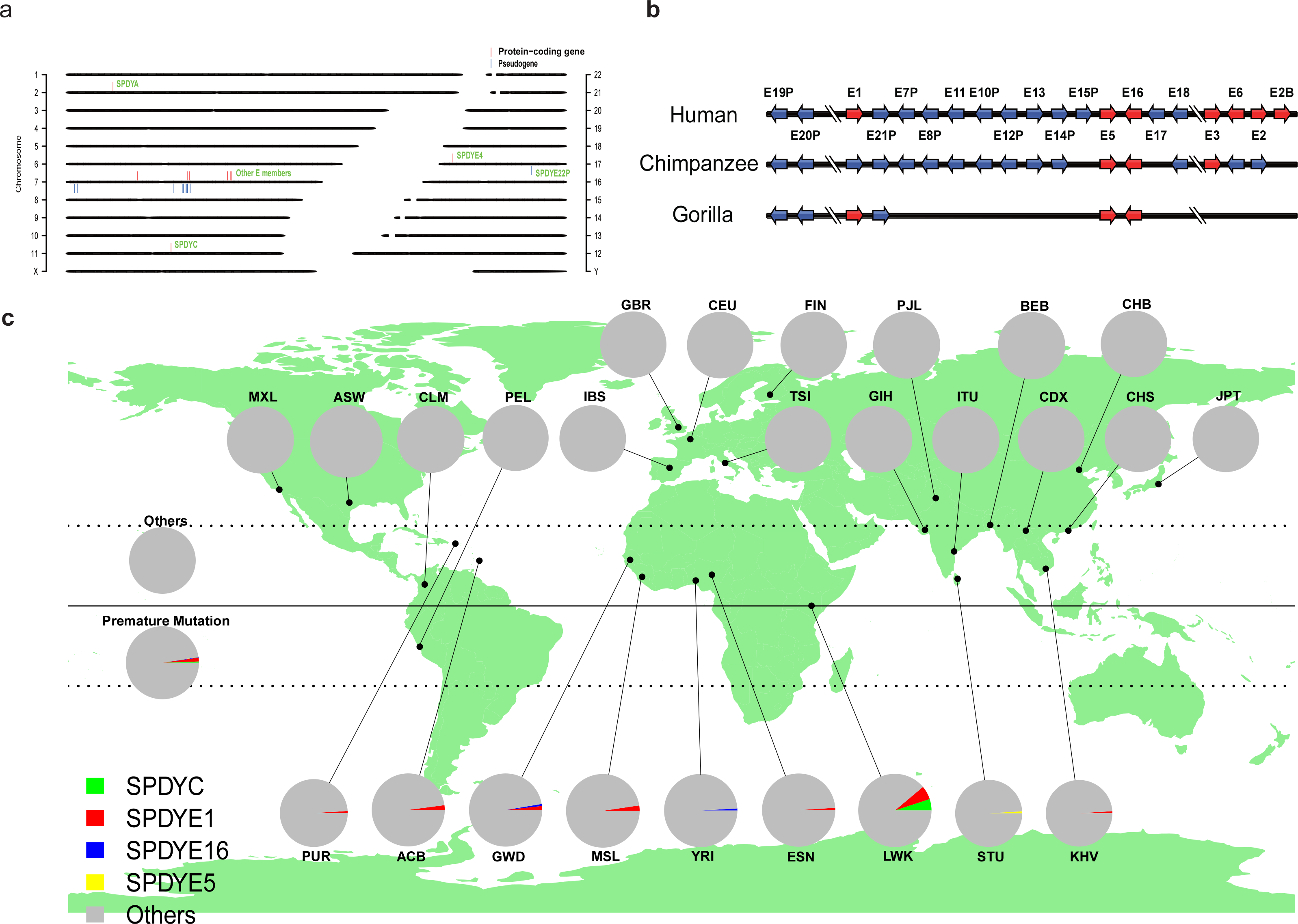
Fixation status of *Speedy* genes in both the Homininae and human populations. (a) The location of *Speedy* genes in the human genome. Red and blue lines represent protein-coding genes and pseudogenes, respectively. Lines above or below the chromosome represent genes that are on the forward or reverse strand, respectively. (b) Schematic comparison of the existence of *Speedy* genes on chromosome 7 among the Homininae. Red and blue arrows represent protein-coding genes and pseudogenes, respectively. Right and left arrows represent forward and reverse strand, respectively. (c) Fixation status of *Speedy* members in 26 human populations. The name of each pie section is assigned based on abbreviation of population name (57). The size of each pie section is proportional to the median of the number of SNPs of individuals in each population. The corresponding proportion of unfixed *Speedy* genes in each population is indicated by the area of different colors.

As the *Speedy* gene subfamily E is not conserved in Homininae, we investigated whether these members are fixed in human populations by using the 1000 Genomes Project data. First, we focused on copy number variations but found no copy number variations in any of the *Speedy* members in 26 human populations. Second, we examined single nucleotide polymorphisms (SNPs) that result in premature termination of CDSs in *Speedy* members. For the pseudogenes, there are SNPs in some individuals that would result in changes from a stop codon to an amino acid, but there are also other stop codons in the sequence (Table S3). We found SNPs that could result in premature termination of CDSs in 4 out of 10 protein-coding genes (Fig. 2c). Most of them came from populations that have more variation sites among genomes (*e.g.*, LWK, ACB, GWD, MSL, ESN, and YRI). Additionally, some members in these populations have a high degree of differentiation, when compared to other populations, especially *SPDYE20P* and *SPDYE15P*. All of the abovementioned populations came from Africa, implicating a strong continental bias for such mutations. The SNPs in the *Speedy*-unfixed equatorial populations were significantly higher than in the other populations (*Mann-Whitney U* test with P<0.001).Furthermore, we found that two sites in *SPDYC* and *SPDYE1* had a high SNP frequency (> 5%) in each sub-population, but only one homozygote for each site. These data indicate that all pseudogenes have been fixed in human populations but protein-coding genes have not.

### Expression and regulation in human normal tissues

Due to the extensive expansion of the *Speedy* family in Catarrhini and its different fixation status in human populations, we assessed whether these genes are expressed in humans. We used transcriptomic data from 32 normal tissues from the Human Protein Atlas to examine their expression profiles. *SPDYE14P* is the only member whose expression could not be detected in any of these tissues. The expression profiles revealed that most *Speedy* members are mainly expressed in the testis (Fig. 3a). We found that *SPDYE11* and *SPDYE4* display significantly tissue-specific expression (tissue-specific value ≥ 0.8), while five other genes (*SPDYE2*, *SPDYE2B*, *SPDYE3*, *SPDYE6*, and *SPDYE12P*) were broadly expressed among normal tissues (with tissue-specific values < 0.1) with similar expression profiles. We also found that *SPDYA*, *SPDYC* and E members have different expression patterns. First, *SPDYA* was not clustered with any other *Speedy* members, indicating its unique expression pattern. Second, in addition to testis, *SPDYC* genes were also expressed in the liver, small intestine, duodenum and placenta. Most E members were highly expressed in the testis except for *SPDYE19P*, *SPDYE11*, and *SPDYE13P*, which also had diverse expression patterns in other normal tissues. Many of them were expressed in the spleen, bone marrow, skin, placenta, stomach, appendix, and lymph node. Although *SPDYE19P*, *SPDYE11* and *SPDYE13P* are not expressed in the testis, they are expressed in other tissues with no obvious rules.

**Figure 3.**
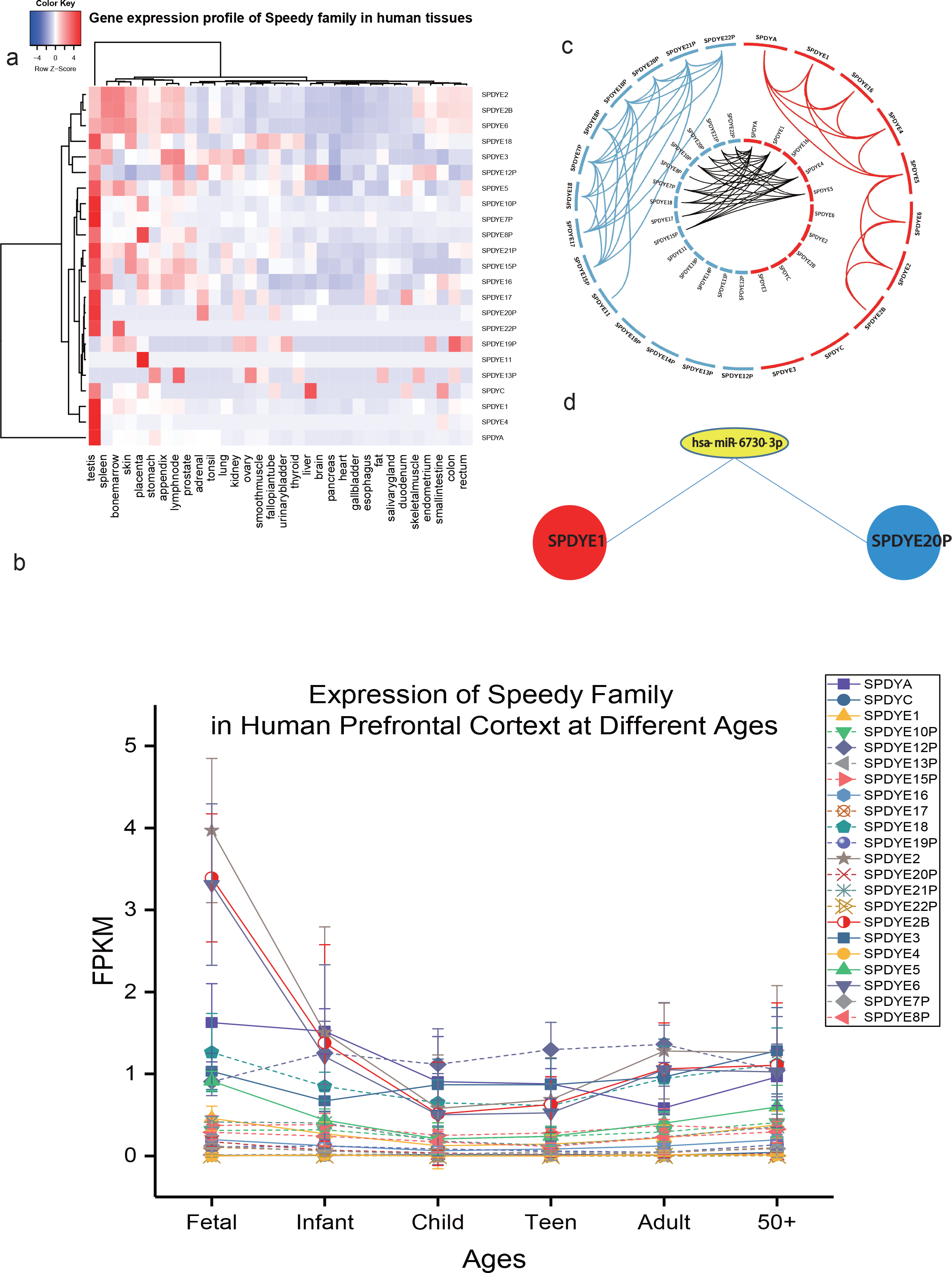
Expression profiles and regulation of *Speedy* members in human normal tissues. (a) Heatmap representation of the expression of *Speedy* members across 32 human tissues. (b) Expression of Speedy members in the human prefrontal cortex at different ages. There are six samples of each age. Solid and dashed lines represent protein-coding genes and pseudogenes, respectively. The median FPKM (fragments per kilobase million) is used for the plot, and standard deviation is displayed by error bars. (c) Circos representation of the expression correlation between *Speedy* members (R^2^≥0.8). The outer circos shows the relationships between protein-coding gene pairs (red) and pseudogene pairs (blue). The inner circos shows gene-pseudogene pairs (black). (d) ceRNA network mediated by a miRNA (has-miR-6730-3p) between *SPDYE1* and *SPDYE20P*.

As new genes have been suggested to drive the development of the human brain, we examined the expression of *Speedy* genes in the human prefrontal cortex. The majority of *Speedy* genes were expressed in the prefrontal cortex (Fig. 3b). In contrast, *SPDYE2, SPDYE2B*, and *SPDYE6* from LSI-Cs had their highest expression levels in stages of fetal development and were decreased across other stages of life, indicating that they may play important roles in early brain development (Fig. 3b). The constitutive expression (median of FPKM ~1) of two protein-coding genes (*SPDYA* and *SPDYE3*) and two pseudogenes (*SPDYE18* and *SPDYE12P*) in the prefrontal cortex across the entire lifespan indicates that they might participate in fundamental functions in the prefrontal cortex. Together, these data demonstrate that *Speedy* genes may have diverse functions, especially in the development of the prefrontal cortex in fetuses.

To determine the potential regulatory relationships among *Speedy* members, we first calculated the correlation coefficient of the expression profile between each gene with the *p* value adjusted by Bonferroni correction. We found that there are significant positive expression correlations among *Speedy* family members (Fig. 3c). We then wondered whether this relationship was the result of competing endogenous RNA (ceRNA) networks, which are mediated by common microRNAs (miRNAs). Five hundred forty-nine miRNAs are predicted (by all 3 methods TargetScan[46], PITA[47] and miRanda[48]) to target *Speedy* members, of which 190 miRNAs belong to 88 known microRNA families. Together with the expression correlation, nine pairs display a ceRNA relationship (Fig. S2). Within ceRNA networks, protein-coding genes and pseudogenes regulate each other. There were also many significant correlations between protein-coding genes and pseudogenes that do not show crosstalk via common miRNAs.

Combined with the origin of pseudogenes, we found that there were two modes of correlation between pseudogenes and their parental genes. First, the expression profile of pseudogenes correlates with their parental genes (*e.g.*, *SPDYE1* and *SPDYE16* with their descendant pseudogenes). Furthermore, *SPDYE20P* and its parental gene *SPDYE1* also share the miRNA target of has-miR-6730-3p (Fig. 3d), indicating that they could regulate each other via ceRNA. Second, the expression profile of pseudogenes does not correlate with their parental genes (*e.g.*, *SPDYE3* and *SPDYE6* with their descendant pseudogenes). Moreover, pseudogenes (*e.g. SPDYE8P*, *SPDYE10P*, *SPDYE15*, and *SPDYE17*) from *SPDYE3* display an expression correlation with other cognates, and *SPDYE19P* from *SPDYE6* does not have any correlation with other members of the *Speedy* family (Fig. 3c and S2).

Since genes in the same pathways or in the same functional complex often exhibit similar expression patterns under diverse temporal or physiological conditions, we examined their possible function(s) by determining co-expression gene networks. We found three protein-coding genes (*SPDYA1*, *SPDYE1*, and *SPDYE4*) that represented co-expressed gene clusters. The gene ontology (GO) functional enrichment analysis by ClueGO[44] and CluePedia[45] revealed that their co-expressed genes are all enriched in reproduction-related functions (Fig. 4), which is consistent with their expression profiles. *SPDYE1* and *SPDYE4* but not *SPDYA* are co-expressed genes with two additional GO terms (*e.g.* phototransduction and acrosome reaction), indicating their functional divergence compared to *SPDYA*.

**Figure 4.**
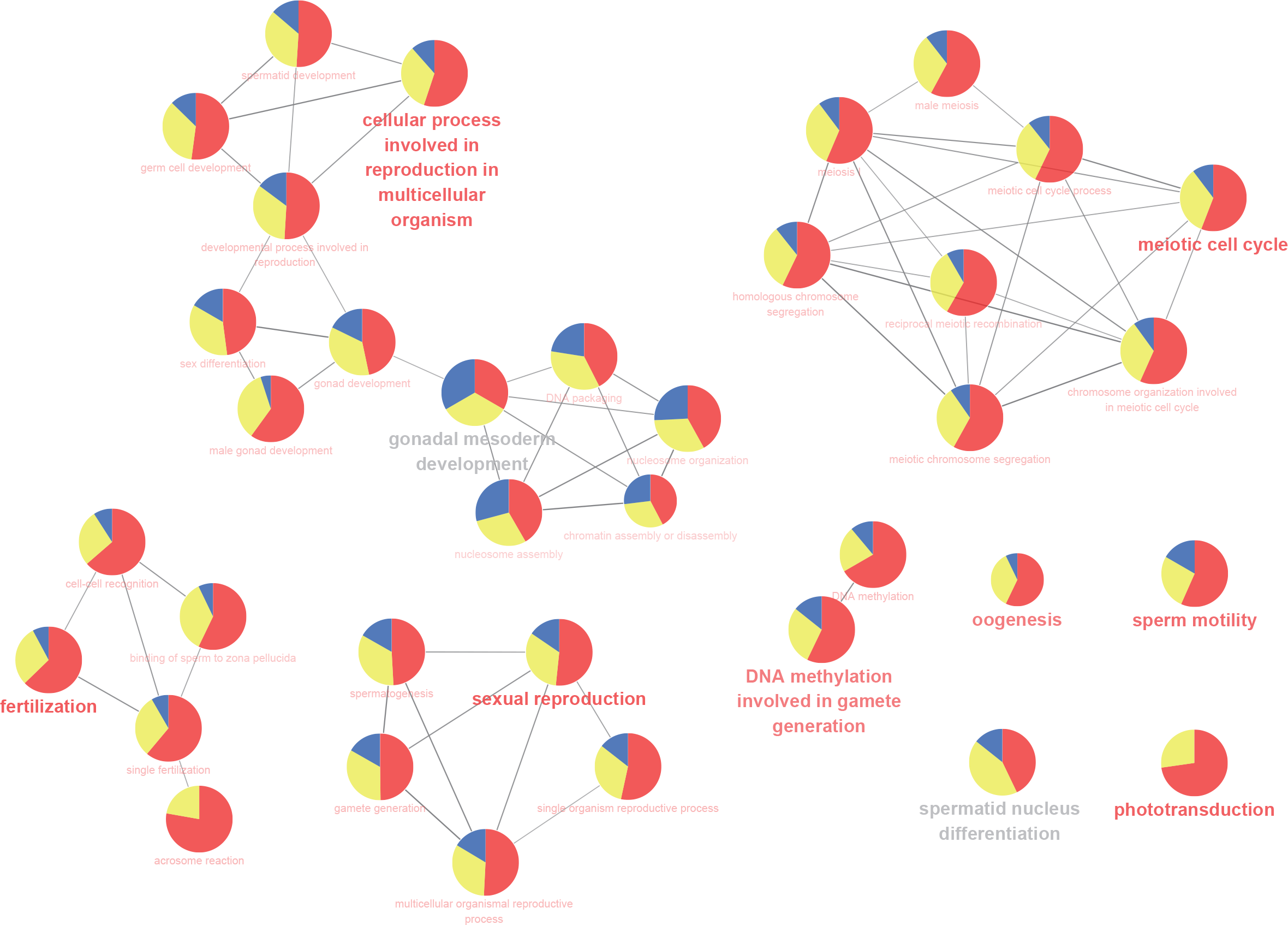
The potential function of *SPDYA*, *SPDYE1* and *SPDYE4* reveals by co-expression networks analysis. Network representation of GO enrichment of genes that are co-expressed with *SPDYA*, *SPDYE1* and *SPDYE4* in human tissues (p<0.05). The color gradient shows the gene proportion of each cluster associated with the term. Node names are assigned according to GO terms. The size of the nodes reflected the statistical significance of the terms (term p-value corrected with Bonferroni). Edges represent the existence of shared genes. The most significant GO term of the group with the lowest term *p*-value is shown in bold using the corresponding color. Equal proportion of each two clusters are represented in grey font.

### *Speedy* family genes are associated with cancer

As *Speedy* genes are expressed in normal human tissues, we next examined whether they are also expressed in tumors. In 734 cancer cell lines (15 cancer types) from the Cancer Cell Line Encyclopedia (CCLE), most pseudogenes are expressed at a lower level in contrast to protein-coding genes (Fig. 5a). However, they do have similar expression profiles, which is consistent with results from normal tissues (Fig. 5b). We only uncovered co-expression gene networks for *Speedy* members from bladder urothelial carcinoma (BLCA) and head and neck squamous cell carcinoma (HNSC). *SPDYE4* was the only *Speedy* member with a co-expression gene network in both normal tissues and HNSC, indicating its fundamental function (Fig. S3).

**Figure 5.**
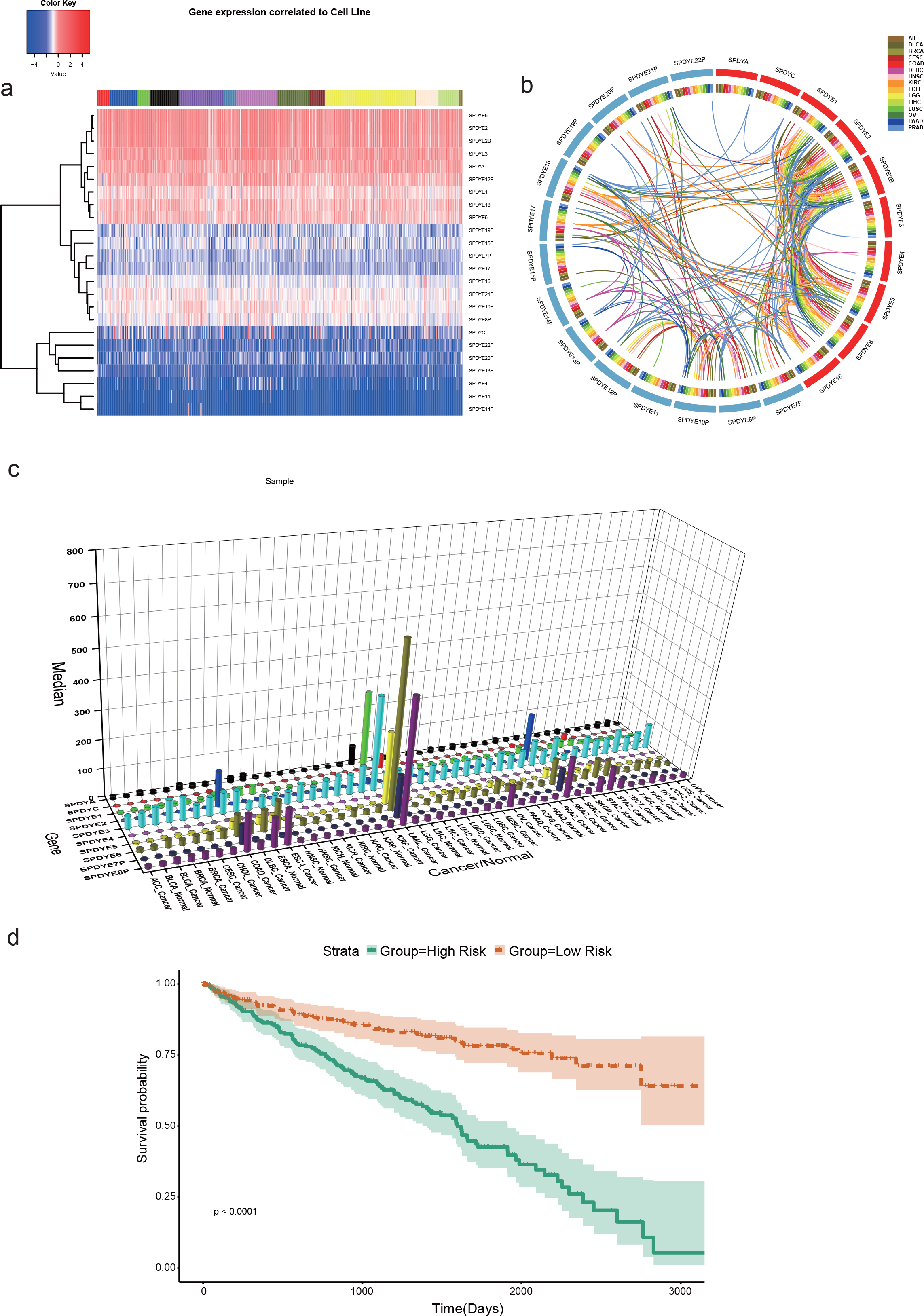
Expression and regulation of *Speedy* members in tumors. (a) Heatmap representation of the expression of *Speedy* members in 734 cancer cell lines. The top color bar represents the 15 cancer types. (b) Circos representation of the expression correlation between *Speedy* members in each cancer type (R^2^≥0.8). Colors represent the expression correlation of two members in different cancer types. (c) Expression profile of *Speedy* members from TCGA. The median of normalized read counts is shown for each gene. (d) Kaplan-Meier analysis of overall survival for patients in KIRC based on the expression signature of five *Speedy* members. Patients were divided into two groups based on the median risk score (0.553\**SPDYA* + 0.195\**SPDYE4* + 0.314\**SPDYE6* - 0.516\**SPDYE7P* + 0.364\**SPDYE8P*). Red and black lines represented high and low risk, respectively. 95% confidence intervals are represented as shaded regions.

In addition to cancer cell lines, we further examined whether the *Speedy* members are expressed in human tumors by using The Cancer Genome Atlas (TCGA) data. Only 10 *Speedy* members were found in TCGA data. Eight out of 10 members have a higher expression level in acute myeloid leukemia (LAML) compared to other cancer types (Fig. 5c). Additionally, *SPDYE2* and *SPDYE7P* also have a high expression level in colon adenocarcinoma (COAD) and rectum adenocarcinoma (READ) compared to other cancer types. In paired data, we found that most members were significantly more highly expressed in tumors than corresponding normal tissues (Table S4). In contrast, *SPDYC* was significantly down regulated in three types of kidney cancer and liver hepatocellular carcinoma (LIHC). Five of six significantly differentially expressed genes were down regulated in thyroid carcinoma (THCA). In kidney renal clear cell carcinoma (KIRC), nine out of 10 members (except for *SPDYE4*) were all differentially expressed between tumors and their corresponding normal tissues.

We further examined whether these 10 *Speedy* members were associated with patient survival in 14 cancer types and found that their expression levels were predictive of patient survival. Especially in KIRC, a high expression level of seven *Speedy* members was all significantly associated with poor patient outcome. We also found that a combination of five *Speedy* members was significantly highly associated with overall survival in KIRC (Fig. 5d). Although *SPDYE4* did not display a significant expression difference between tumors and corresponding normal tissues in KIRC, prostate adenocarcinoma (PRAD) and skin cutaneous melanoma (SKCM), its expression level was able to predict patient survival (Table S5). In addition to protein-coding genes, *SPDYE7P* and *SPDYE8P* were also significantly associated with patient survival in KIRC/SKCM and HNSC/KIRC/Mesothelioma (MESO), respectively. Notably, *SPDYA* had an inverse relationship with patient survival among different cancer types (*e.g.*, a higher expression level with longer survival in LAML and lung squamous cell carcinoma (LUSC) and a lower expression level with poor survival in KIRC and PRAD). Besides, human specific *Speedy* genes (*SPDYE2* and *SPDYE6*) were also linked to cancer. *SPDYE2* was upregulated in BRCA, KICH, KIRC, KIRP, LUAD, and LUSC, while *SPDYE6* was upregulated in BLCA, BRCA, KIRC, LIHC, LUSC, and downregulated in THCA. High expression level of *SPDYE2* was associated with the poor patient survival in KIRC, SKCM, and UVM. High expression level of *SPDYE6* was associated with the poor patient survival in KIRC, and UVM. However, low expression level of *SPDYE6* was associated with the poor patient survival in DLBC (Fig. S4).

Overall, members of the *Speedy* family display different expression profiles and different associations with patient survival in different cancer types. This indicates that they may be related to cancer to some extent and are potential biomarkers for different cancer types.

## Discussion

In this study, we comprehensively identified *Speedy* genes. Our analysis and the published eukaryotic species tree [51] revealed that *Speedy* is a very old gene. Although there are two species in Amoebozoa containing *Speedy* genes, it can still be concluded that the *Speedy* gene originated at latest before the common ancestor of the Metazoa. This family was lost in many species, with no obvious taxonomic rules during evolution. The absence of the *Speedy* family could notbe completely attributed to incomplete genome sequences and/or annotations, *e.g.*, fruit fly is well annotated, but we didn’t find any *Speedy* genes in it. In addition to the existence bias across species, the number of *Speedy* genes varied among different species. However, it is obvious that higher eukaryotes have more *Speedy* genes than lower ones. The uneven distribution of the *Speedy* family is also a functional hint that *Speedy* gene may be play roles in some subtle regulatory processes instead of being indispensable for life. In this study, we used the comprehensive sequence dataset, so that the *Speedy* gene family could be further spilt into three subfamilies instead of two. As there were only two subfamily C genes (*Speedy* B4-like) in the previous study [14], it would be possible that they were not distinguishable from the *Speedy* B subfamily [14] (corresponding to the *Speedy* E subfamily in this study). To clarify the situation, we followed the official gene names of the human *Speedy* genes approved by the HUGO Gene Nomenclature Committee and split the previous *Speedy* B subfamily genes [14] into the *Speedy* C and *Speedy* E subfamilies.

Gene duplication during evolution is one genetic source of gene family expansion, by means of acquiring genetic novelty in organisms [52]. When a paralog arises in an individual genome, it follows three evolutionary stages to maintain themselves in a population: fixation, fate-determination, and preservation[53]. In our study, a novel domain architecture was observed in *Speedy* subfamily E, and the poor-identity-*spy1* domain is under negative selection, indicating that it has a potential unknown function. Twenty-four human *Speedy* members were identified in our study. Among them, 10 are protein-coding genes, of which *SPDYE5* and *SPDYE16* also exist in both chimpanzees and gorillas with coding potential. This suggests that these two genes were already be functional in the common ancestor of the Homininae. Fourteen pseudogenes were also found in the human genome, which were all paralogs LSI-Cs and the subsequent mutation accumulation of premature stop codons. When compared to the chimpanzee and gorilla genomes, we found that all human pseudogenes are pseudogenized in chimpanzees, but most do not exist in gorilla genome. The majority of protein-coding genes in the *Speedy* subfamily E also display similar trends. The massive extinction of members of subfamily E in gorillas could be an artifact of the low quality of its genome sequences[54]. Alternatively, it could indicate that expanded members from subfamily E are not conserved among the Homininae. When we examined the fixation status of *Speedy* members at the human population level, we found that pseudogenes are already fixed in human populations, but some protein coding genes are not completely fixed in several equatorial populations. These populations, *per se*, have a higher mutation rate than others who reside beyond the equator. The higher mutation rate in these population would account for the non-fixation status of some *Speedy* members.

After fixation, paralogs can face different destinies, such as pseudogenization, retaining the same function as parental genes, subfunctionalization, or neofunctionalization, *etc*[55]. Such new genes, especially lineage-specific ones, are also common targets of positive selection[56] and therefore could be very important in adaptation[57]. We did observe that the *Speedy* subfamily E in Catarrhini experienced positive selection. In addition, type ◻ divergence analysis indicated that all proteins of subfamily E have different functions compared to subfamily A and C. *Speedy* E members also displayed different binding preferences for CDKs and effects on the cell cycle. Taken together, the data suggest that the newly evolved subfamily E genes display functional diversity due to positive selection.

In addition to information from the sequence level, gene expression profiles also more directly shed light on their functional status. Most members of human *Speedy* subfamily E are mainly expressed in the testis, and some are also expressed in the brain, which is consistent with previous studies demonstrating that new genes are often predominantly expressed in the testis and human brain[57, 58]. Although the testis-biased expression could be the result of the widespread transcriptional activity in the testis[59], genes that are mainly expressed during gametogenesis or have an effect on reproduction often undergo positive selection[60], and new genes could potentially create reproductive barriers. In this case, *Speedy* members experienced positive selection and displayed significant reproduction-related functions. Thus, they may play an important role in reproductive barriers. *SPDYE4* is mainly expressed in testis and further functional enrichment analysis of genes that are co-expressed with *SPDYE4* suggesting that *SPDYE4* might play an important role in meiosis.

It is well known that the prefrontal cortex, which is in charge of cognitive abilities, expanded in size and complexity in primates [61]. *SPDYE2* displayed a higher expression level in the fetal prefrontal cortex than during other periods, indicating *SPDYE2* plays an essential role in the development of the prefrontal cortex. According to the expression profiles of other *Speedy* members in the prefrontal cortex at different ages, we have no reason to doubt that *Speedy* genes are involved in cognitive ability to some extent.

Given that paralogous genes have similar or even identical expression patterns, it is suggested that they could be functionally identical or redundant [62]. We found that most protein coding genes and pseudogenes have identical expression profiles as their counterparts, implicating functional redundancy. The diversity of expression profiles between *Speedy* members also suggests that they may have different functions in different tissues. Taken together, when we focused on human *Speedy* subfamily E genes, their expression profiles might be explained by the out-of-testis hypothesis that new genes are apt to initially gain function in testis and then extend their expression into other tissues to acquire new functions[63, 64]. Additionally, significant expression correlations were found between *Speedy* genes and pseudogenes and some of them shared common miRNA targets.

In 2011, the ceRNA hypothesis was proposed, which describes a novel and complicated post-transcriptional regulation network containing transcribed pseudogenes. mRNAs and transcribed pseudogenes are connected by common miRNAs[65]. This provide new perspective to examine the function of transcribed pseudogenes. Other studies show that mRNA can act as both a protein coding sequence and ceRNA [66]. It is assumed that even if a protein-coding gene lost its coding capacity (pseudogenization), it would retain its ceRNA function if expressed [40].

Paralogous genes may suffer from dosage balance and co-regulation between each other[67]. For *Speedy* members, we also observed the phenomenon that most pseudogenes are expressed at low levels in normal tissues. Additionally, we identified some pseudogenes that share common miRNAs with protein-coding genes in the *Speedy* family. Combined with the results of expression profile correlation, the origins of pseudogenes, and sharing miRNAs targets, we proposed two major regulatory modes. One is that pseudogenes can regulate their parental genes or others by their transcripts acting as ceRNAs; the other is that pseudogenes can only regulate cognates of their parental genes. Phylogenetic analyses suggest that *SPDYE3* is the ancestor of human LSI-Cs, but the expression of pseudogenes originating from *SPDYE3* does not display correlation with *SPDYE3* expression. It is obvious that *SPDYE3* is an ancestor of LSI-Cs and generated many ‘descendants’, among which some became pseudogenes that then regulated their ‘brothers’. This process also generated many pseudogenes that regulate themselves. Human *Speedy* family members thus formed a ‘regulatory flux’ during evolution. Human *Speedy* members also comprise a ceRNA network to regulate each other after extensive duplication. The ceRNA networks that we reconstruct are underestimated, due to the strict criteria of expression profile correlation and miRNA target prediction.

*Speedy* members are mainly expressed in the testis, and the processes of gametogenesis and tumorigenesis share important similarities, indicating that *Speedy* genes may also play an important role in tumors. We did detect their expression in cancer cell lines and tumors. Among 32 cancer types, eight out of 10 *Speedy* members were highly expressed in LAML, except for *SPDYC* and *SPDYE2*. This is a clue that *Speedy* genes might be related to cancer, especially in LAML. From normal tissues to cancer cell lines and tumor samples, we found that *SPDYE14* were not expressed or had a very low expression level, indicating that it is likely a non-functional pseudogene. *SPDYE4* is a protein-coding gene and displayed restrictedly expression in the testis and cancer, indicating it as a potential cancer/testis (CT) antigen (a class of tumor antigens with testis-specific expression in human normal tissues, or in some cases such as ovary and trophoblast)[68]. Due to the characteristics of its expression profile, *SPDYE4* is a potential therapeutic target for cancer. We also found that *Speedy* members are significantly associated with overall survival for patients in diverse cancer types. However, the expression level could be either associated with better or worse overall survival for patients depending on cancer type. Together with the different expression profiles between tumor and normal tissues, we speculate that *Speedy* members can be used as novel biomarkers and therapeutic targets in different cancer types. The expression characteristics of E members in normal and cancer cells show that they could bring both advantageous function and negative effects to organisms, which is consistent with the pleiotropy of new genes. Taken together with the positive selection on E members, it might be a consequence of further evolution to ‘solve’ a newly negative problems brought about by the fixation of new genes as compensatory changes, which is the derivation of selection, pleiotropy, and compensation hypothesis [64, 69].

## Conclusions

In general, our study shows that *Speedy* family genes are extensively expanded in the Homininae via the formation of a novel subfamily containing a low-Spy1-identity domain. Although they are still not completely fixed among Homininae and human populations after expansion, their expression profiles show that they have two distinct regulatory modes and play an important role in human brain. We also demonstrated their clinical relevance in diverse cancer types and imply that they are potential novel biomarkers and therapeutic targets for cancer.

## Abbreviations

ceRNA: competitive endogenous RNA
FPKM: Fragments Per Kilobase of transcript per Million mapped reads
LSI-C: *Speedy* gene with low-identity *Spy1* domain located at C-terminal
CPV: coding potential value
CDS: coding DNA sequence
SNP: single nucleotide polymorphism
GO: gene ontology
CCLE: Cancer Cell Line Encyclopedia
BLCA: bladder urothelial carcinoma
HNSC: head and neck squamous cell carcinoma
TCGA: The Cancer Genome Atlas
LAML: acute myeloid leukemia
COAD: colon adenocarcinoma
READ: rectum adenocarcinoma
LIHC: liver hepatocellular carcinoma
THCA: thyroid carcinoma
KIRC: kidney renal clear cell carcinoma
PRAD: prostate adenocarcinomaSKCM: skin cutaneous melanoma
SKCM: skin cutaneous melanoma
MESO: Mesothelioma
LUSC: lung squamous cell carcinoma

## Declarations

### Ethics approval and consent to participate

Not applicable.

### Consent for publication

Not applicable.

### Availability of data and material

The human genome variation datasets analyzed during the current study are available in the 1000 Genomes Project website [http://www.internationalgenome.org]. RNA-Seq data of cancer cell lines, analyzed during the current study are available at NCBI Gene Expression Omnibus with accession number GSE36139. RNA-Seq data of human normal tissues and human prefrontal cortex Sequence Read Archive database under bioproject number PRJNA183192 and PRJNA245228, respectively. The datasets of *Speedy* gene sequences generated and analyzed during the current study are available from the corresponding author on reasonable request.

### Authors’ contributions

S.G., G.F.G. and L.W. designed and supervised the study. L.W. and H.W. analyzed data. L.W. Hongmei, W., H.W., Y.Z., X.L., Q.Y., X.X., G.F.G. and S.G. wrote the paper. And L.W. Hongmei, W., H.W., Y.Z., X.L., G.W, Q.Y., X.X., G.F.G. and S.G. revised the paper.

## Acknowledgments

We thank Manyuan Long from the University of Chicago, Wen Wang from the Kunming Institute of Zoology, Chinese Academy of Science, and Zhijin Liu from the Institute of Zoology, Chinese Academy of Science for helpful advice and discussions.

